# Specific auxin and medium combinations alter *Saccharomyces cerevisiae* growth

**DOI:** 10.1101/2025.09.17.674911

**Authors:** Neta Nnabuenyi, Morgan A Sands, Nicole J Camlin

## Abstract

Auxin-inducible degradation is an increasingly popular protein-targeting reverse genetics approach. Use of this method has revolutionized the types of questions cell and molecular biologists can answer, however, a growing number of studies point to auxin alone impacting different cellular phenotypes. This study investigated the impact of different medium and auxin combinations on *Saccharomyces cerevisiae* growth. We observed that both natural and synthetic auxin (Indole-3-acetic acid (IAA) and 1-Naphthaleneacetic acid (NAA) respectively) impaired budding yeast growth in nutrient minimal but not nutrient rich media. This finding was true across different yeast strains with or without an intact auxin-inducible degradation system. Ultimately, this study highlights the need for proper controls when using auxin-inducible degradation.

## Description

Protein-targeting reverse genetic approaches are becoming increasingly popular. Auxin-inducible degradation represents one of the most widespread methodologies to quickly and temporally remove proteins-of-interest (POI). This system has three parts. First, the plant F-box TIR1 (from *Oryza sativa* (osTIR1) or from *Arabidopsis thaliana* (atTIR1)) is exogenously expressed in non-plant eukaryotic cells. TIR1 is able to form a functional SCF E3 ligase with endogenous proteins (Figure 1A). Secondly, POI are tagged with the TIR1 recognition motif, auxin-inducible degron (AID). Finally, the plant hormone auxin is added to the system. This induces TIR1-AID-tagged POI interaction leading to polyubiquitination and proteasomal degradation of the POI (Nishimura et al., 2009). Use of this system is becoming increasingly widespread in diverse fields of cell biology and across model organisms (Brown et al., 2017; Camlin & Evans, 2019; Holland et al., 2012; Kanke et al., 2011; Kreidenweiss et al., 2013; Nishimura et al., 2009; Zhang et al., 2015). As a result, auxin-inducible degradation has led to significant advancements in our understanding of cellular functions. However, like all methodological approaches, this system is not without its drawbacks. A growing number of papers have found that auxin is not as inert to non-plant eukaryotic cells as initially believed. For example, in *Drosophila* auxin was found to alter gene expression (Fleck et al., 2024). Furthermore, in *Saccharomyces cerevisiae*, the naturally occurring auxin Indole-3-acetic acid (IAA) has been found to impair cell growth, alter the TORC1 pathway, induce filamentation, and impair mRNA movement (Domeni Zali & Moriel-Carretero, 2023; Nicastro et al., 2021; Prusty et al., 2004).

**Figure Legend:**
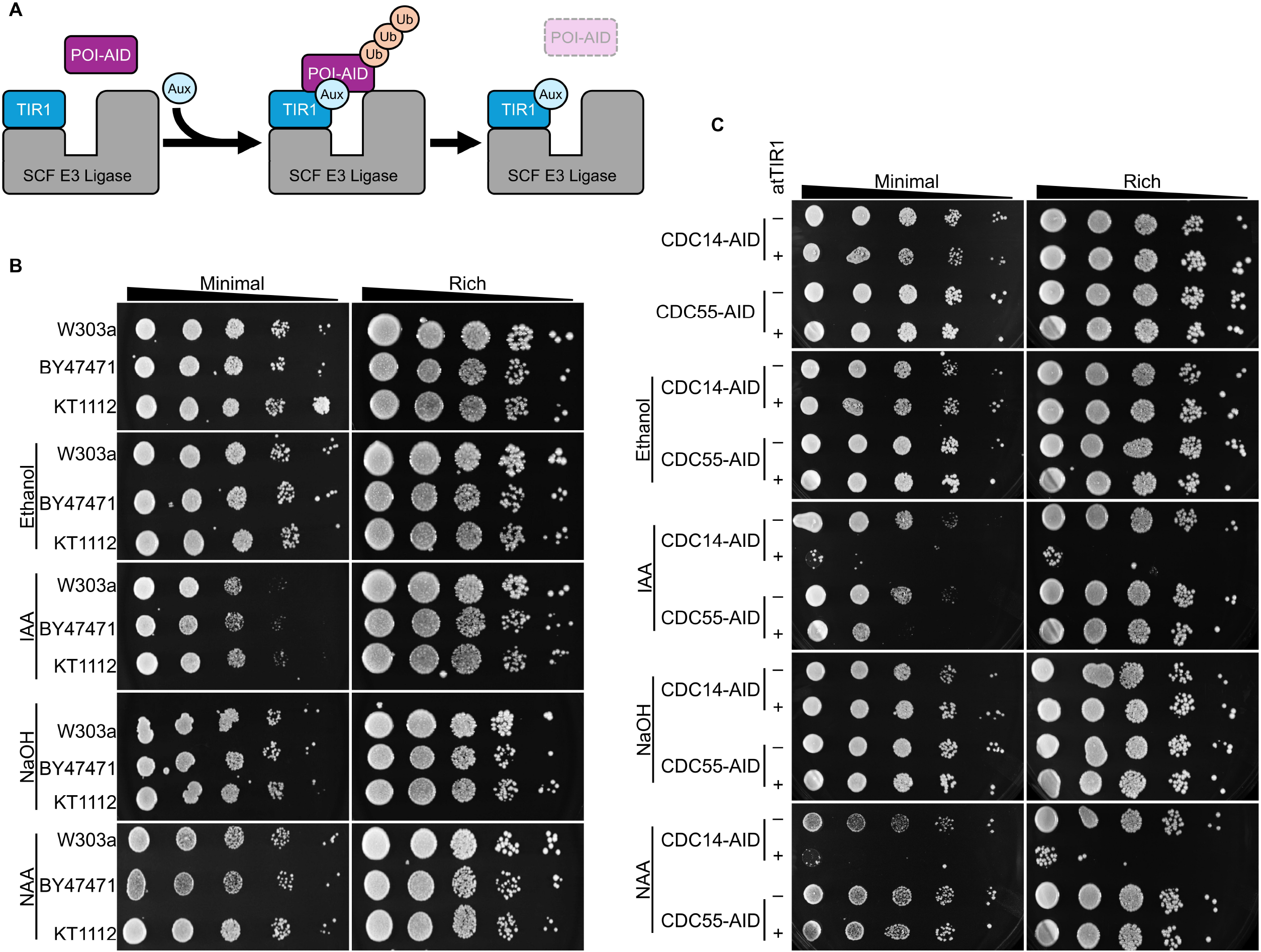
Impact of auxin on different *S. cerevisiae* strains. (A) Systematic representation of auxin-inducible degradation. Exogenous TIR1 is expressed in non-plant eukaryotic cells. TIR1 forms a functional E3 ligase with endogenous SCF proteins. The protein-of-interest (POI) is tagged with the auxin-inducible degron (AID). Under standard culture conditions TIR1 and POI-AID do not interact. Upon the addition of auxin (Aux) TIR1 interacts with POI-AID leading to polyubiquitination (Ub) and ultimately proteasomal degradation of AID-tagged POI. (B) Growth spotting assay for three different wild type *S. cerevisiae* strains grown for 48 hours on minimal or rich media. Plates were supplemented with ethanol (IAA vehicle), IAA, NaOH (NAA vehicle), NAA or nothing (control plate). (C) Growth spotting assay for yeast with an AID-tagged POI, CDC14 or CDC55, with and without atTIR1. *S. cerevisiae* was grown for 48 hours on minimal or rich media supplemented with ethanol, IAA, NaOH, NAA, or nothing.

While there is precedent for auxin impacting *S. cerevisiae* growth, to date no systematic assessment has been performed. Recently our laboratory noticed a previously unpublished phenomenon, variable *S. cerevisiae* growth with specific auxin and medium combinations. Therefore, we set out to systematically test this phenotype across multiple budding yeast strains to determine if this was a universal impact across multiple yeast strains or if this was specific to certain backgrounds. Experiments compared the impact of different auxin types (IAA or 1-Naphthaleneacetic acid (NAA)) on cell growth in nutrient rich (YPD medium) or minimal (SCD medium) conditions (Figure 1B). Initial experiments compared three commonly used wildtype yeast strains W303a, BY4741, and KT1112. Auxin (500 µM IAA or NAA) or their vehicle controls (ethanol or NaOH) had no impact on yeast growth compared to control plates (no auxin or vehicle) in nutrient rich medium. Conversely, both IAA and NAA reduced yeast growth on minimal media plates for all wild type strains tested.

To determine if this phenotype was also observed in *S. cerevisiae* with a functional auxin-inducible degradation system, a second set of experiments were conducted using yeast strains with AID-tagged endogenous proteins with and without integrated atTIR1 (Figure 1C). Of note, an essential (*Cdc14*) and non-essential (*Cdc55*) gene was chosen for this assay *(Powers & Hall, 2017; Visintin et al., 1998; Yellman & Burke, 2006)*. As expected, auxin-inducible degradation of CDC14-AID significantly impaired yeast growth as previously observed (Powers & Hall, 2017). Conversely, loss of CDC55-AID led to no observable impact. However, as with wildtype strains, addition of auxin (IAA or NAA) to minimal medium impaired growth in all cells, whether an essential protein was degraded or not. Taken together these results clearly show that choice of auxin medium combination is extremely important for auxin-inducible degradation experiments. Therefore, care should be taken to ensure adequate control for auxin impact is included when using auxin-inducible degradation.

## Methods

*S. cerevisiae* were grown to stationary phase at 30°C in YPD (yeast, peptone, dextrose) medium. Cells were diluted to an OD600 of 1 with sterile water. Ten-fold serial dilutions were spotted onto agar plates (YPD or SCD (synthetic complete + dextrose)) with 500 µM IAA or NAA, vehicle (ethanol for IAA and NaOH for NAA), or with no additive. Plates were grown at 30°C for 48 h with images taken on a ChemiDoc MP Imaging System (BioRad) at 24 and 48 h.

## Reagents

| ***Saccharomyces cerevisiae* strains** |  |  |
| --- | --- | --- |
| **Strain** | **Genotype** | **Source** |
| W303a | MATa ura3-52 trp1Δ2 leu2-3_112 his3-11 ade2-1 can1-100 | Horizon Discovery - YSC1058 |
| BY4741 | MATa his3Δ1 leu2Δ0 met15Δ0 ura3Δ | Horizon Discovery - YSC1048 |
| KT1112 | MATα ura-52 leu2 his3 | Kelly Tatchell (Stuart et al., 1994) |
| YKA890 | W303 MATa Cdc14-V5/AID:KanMX | Mark Hall (Powers & Hall, 2017) |
| YKA892 | W303 MATa Cdc14-V5/AID:KanMX leu2::ADH1p-AtTIR1::LEU2 | Mark Hall (Powers & Hall, 2017) |
| YKA1223 | W303 MATa CDC55-V5/AID:NAT | Mark Hall |
| YKA1229 | W303 MATa CDC55-V5/AID:NAT trp1::ADH1p-AtTIR1(F79A):TRP1 | Mark Hall |

| Medium |  |  |
| --- | --- | --- |
| Reagent | Source | Catalogue Number |
| Yeast Extract | Fisher Scientific | BP1422 |
| Peptone | Fisher Scientific | BP1420 |
| Dextrose | Fisher Scientific | D16 |
| Synthetic complete amino acid supplement w/o Yeast Nitrogen Base | USBiological | D9515 |
| Yeast Nitrogen Base without Amino Acids | Research Products International Corp | Y20040 |
| 1-Naphthaleneacetic acid | Sigma Aldrich | N0640 |
| Indole-3-acetic acid | Gold Biotechnology | I-110-25 |
| Agar, Powder | Fisher Scientific | BP1423 |

## Acknowledgements

The authors greatly acknowledge the yeast strains provided by Kelly Tatchell (Louisiana State University Health Sciences Center) and Mark Hall (Purdue University).

## Author Contributions

Neta Nnabuenyi: Investigation, Writing - review and editing

Morgan A Sands: Investigation, Funding acquisition, Writing - review and editing

Nicole J Camlin: Conceptualization, Methodology, Project administration, Investigation, Supervision, Funding acquisition, Writing - original draft, Writing - review and editing

## References

Brown, K. M., Long, S., & Sibley, L. D. (2017). Plasma Membrane Association by N-Acylation Governs PKG Function in Toxoplasma gondii. mBio, 8(3). 10.1128/mBio.00375-17

Camlin, N. J., & Evans, J. P. (2019). Auxin-inducible protein degradation as a novel approach for protein depletion and reverse genetic discoveries in mammalian oocytes. Biology of Reproduction, 101(4), 704–718. 10.1093/biolre/ioz113

Domeni Zali, G., & Moriel-Carretero, M. (2023). Auxin alone provokes retention of ASH1 mRNA in Saccharomyces cerevisiae mother cells. MicroPubl Biol, 2023. 10.17912/micropub.biology.000752

Fleck, S. A., Biswas, P., DeWitt, E. D., Knuteson, R. L., Eisman, R. C., Nemkov, T., D’Alessandro, A., Tennessen, J. M., Rideout, E., & Weaver, L. N. (2024). Auxin exposure disrupts feeding behavior and fatty acid metabolism in adult Drosophila. Elife, 12. 10.7554/eLife.91953

Holland, A. J., Fachinetti, D., Han, J. S., & Cleveland, D. W. (2012). Inducible, reversible system for the rapid and complete degradation of proteins in mammalian cells. Proceedings of the National Academy of Sciences, 109(49), E3350–E3357. 10.1073/pnas.1216880109

Kanke, M., Nishimura, K., Kanemaki, M., Kakimoto, T., Takahashi, T. S., Nakagawa, T., & Masukata, H. (2011). Auxin-inducible protein depletion system in fission yeast. BMC Cell Biology, 12(1), 8. 10.1186/1471-2121-12-8

Kreidenweiss, A., Hopkins, A. V., & Mordmüller, B. (2013). 2A and the Auxin-Based Degron System Facilitate Control of Protein Levels in Plasmodium falciparum. PLOS ONE, 8(11), e78661. 10.1371/journal.pone.0078661

Nicastro, R., Raucci, S., Michel, A. H., Stumpe, M., Garcia Osuna, G. M., Jaquenoud, M., Kornmann, B., & De Virgilio, C. (2021). Indole-3-acetic acid is a physiological inhibitor of TORC1 in yeast. PLoS Genet, 17(3), e1009414. 10.1371/journal.pgen.1009414

Nishimura, K., Fukagawa, T., Takisawa, H., Kakimoto, T., & Kanemaki, M. (2009). An auxin-based degron system for the rapid depletion of proteins in nonplant cells [10.1038/nmeth.1401]. Nat Methods, 6(12), 917–922. 10.1038/nmeth.1401

Powers, B. L., & Hall, M. C. (2017). Re-examining the role of Cdc14 phosphatase in reversal of Cdk phosphorylation during mitotic exit. Journal of Cell Science, 130(16), 2673. 10.1242/jcs.201012

Prusty, R., Grisafi, P., & Fink, G. R. (2004). The plant hormone indoleacetic acid induces invasive growth in Saccharomyces cerevisiae. Proceedings of the National Academy of Sciences, 101(12), 4153–4157. 10.1073/pnas.0400659101

Stuart, J. S., Frederick, D. L., Varner, C. M., & Tatchell, K. (1994). The mutant type 1 protein phosphatase encoded by glc7-1 from Saccharomyces cerevisiae fails to interact productively with the GAC1-encoded regulatory subunit. Mol Cell Biol, 14(2), 896–905. 10.1128/mcb.14.2.896-905.1994

Visintin, R., Craig, K., Hwang, E. S., Prinz, S., Tyers, M., & Amon, A. (1998). The Phosphatase Cdc14 Triggers Mitotic Exit by Reversal of Cdk-Dependent Phosphorylation. Molecular Cell, 2(6), 709–718. 10.1016/S1097-2765(00)80286-5

Yellman, C. M., & Burke, D. J. (2006). The Role of Cdc55 in the Spindle Checkpoint Is through Regulation of Mitotic Exit in Saccharomyces cerevisiae. Molecular Biology of the Cell, 17(2), 658–666. 10.1091/mbc.e05-04-0336

Zhang, L., Ward, J. D., Cheng, Z., & Dernburg, A. F. (2015). The auxin-inducible degradation (AID) system enables versatile conditional protein depletion in C. elegans. Development, 142(24), 4374–4384. 10.1242/dev.129635

